# Despite impaired binocular function, binocular disparity integration across the visual field is spared in normal aging and glaucoma

**DOI:** 10.1101/2022.11.28.518250

**Authors:** Guido Maiello, MiYoung Kwon

## Abstract

**Objective:** To examine how binocularly asymmetric glaucomatous visual field damage affects processing of binocular disparity across the visual field.

**Design:** Case–control study.

**Participants and Controls:** A sample of 18 patients with primary open-angle glaucoma, 16 age-matched controls, and 13 young controls.

**Methods:** Participants underwent standard clinical assessments of binocular visual acuity, binocular contrast sensitivity, stereoacuity, and perimetry. We employed a previously validated psychophysical procedure to measure how sensitivity to binocular disparity varied across spatial frequencies and visual field sectors, i.e. with full-field stimuli spanning the central 21° of the visual field, and with stimuli restricted to annular regions spanning 0°-3°, 3°-9° or 9°-21°.

**Main Outcome Measures:** We verified the presence of binocularly asymmetric glaucomatous visual field damage by comparing—between the two eyes— the mean deviation values obtained from the Humphrey Field Analyzer (HFA) 24-2 test. To assess the spatial-frequency tuning of disparity sensitivity across the visual field of patients and controls, we fit disparity sensitivity data to log-parabola models and compared fitted model parameters. Lastly, we employed disparity sensitivity measurements from restricted visual field conditions to model different possible scenarios regarding how disparity information is combined across visual field sectors. We adjudicated between the potential mechanisms by comparing model predictions to the observed patterns of disparity sensitivity with full-field stimuli.

**Results:** The interocular difference in HFA 24-2 mean deviation was greater in glaucoma patients compared to both young and age-matched controls (ps=.01). Across participant groups foveal regions preferentially processed disparities at finer spatial scales, whereas periphery regions were tuned for coarser scales (p<.001). Disparity sensitivity also decreased from the fovea to the periphery (p<.001) and across participant groups (ps<.01). Finally, similar to controls, glaucoma patients exhibited near-optimal disparity integration, specifically at low spatial frequencies (p<.001).

**Conclusions:** Contrary to the conventional view that glaucoma spares central vision, we find that glaucomatous damage causes a widespread loss of disparity sensitivity across both foveal and peripheral regions. Despite these losses, cortical integration mechanisms appear to be well preserved, suggesting that glaucoma patients make the best possible use of their remaining binocular function.

---

Glaucoma is a leading cause of irreversible blindness worldwide, characterized by progressive loss of retinal ganglion cells and resultant visual field defects^1^. The loss and/or dysfunction of retinal ganglion cells often leads to a detrimental effect on various visual functions ranging from simple light detection to complex everyday tasks such as object/face recognition, visual search, reading, and mobility^2–13^, thereby affecting quality of life. Primary open angle glaucoma, the most common type of glaucoma, is putatively associated with peripheral vision loss. However, there is accumulating evidence that even early glaucomatous injury may involve the macula, and that such macular damage may be more common than generally thought^14–21^. For example, a number of anatomical studies^16,19,20,22–26^ using Spectral-Domain Optical Coherence Tomography have shown that the thickness of the retinal nerve fiber layer and the retinal ganglion cell plus inner plexiform layer, even in the macula, are significantly thinner in patients with early glaucoma than in healthy controls. Furthermore, glaucomatous visual field loss often occurs asymmetrically not only between two eyes^27^, but also across the visual field^28^. This inhomogeneous nature of glaucomatous visual field deficits is assumed to result in the deterioration of binocular function^29^, which in turn impacts the performance of various everyday visual tasks, such as reading, object recognition, and visuomotor coordination^30–32^. Indeed, studies have shown that, even in early or moderate stages, stereopsis, convergence, and binocular fusion are significantly impaired in people with glaucoma compared to glaucoma suspects or normal cohorts^33–35^.

The perception of disparity-defined depth is a hallmark of binocular visual function. Previous studies in young and healthy normal vision have shown that disparity is processed at different spatial scales in different regions of the visual field^36,37^: the fovea preferentially processes disparities at fine spatial scales, whereas the visual periphery is tuned for coarse spatial scales. Since glaucomatous damage may affect both foveal and peripheral visual field regions asymmetrically in the two eyes, it remains unclear how glaucomatous damage affects disparity sensitivity across the visual field. Moreover, to recover the depth structure of the environment, the healthy visual system selects and combines depth information processed throughout the visual field in near-optimal fashion, i.e. by accounting for the relative reliability of the disparity signals coming from different visual field sectors^36^. However, the question arises as to whether disparity information would be combined in such near-optimal manner as shown in normal healthy vision even if disparity signals are asymmetric across the visual field, as expected in glaucomatous vision. In particular, widespread changes in the brain have been shown as a secondary consequence of retinal ganglion cell loss from glaucoma largely due to direct and transsynaptic anterograde axonal degeneration^38–40^. Therefore, it remains to be seen how glaucomatous damage may affect the visual system’s ability to combine depth information throughout the visual field. The objectives of the current study were thus two-fold: (i) to elucidate whether disparity sensitivity is visual-field dependent in glaucomatous vision and (ii) to identify the mechanisms underlying disparity integration across the visual field.

To address the first goal, we measured disparity sensitivity across the visual field of glaucoma patients and healthy controls using annular pink noise stimuli embedded with disparity corrugations of different spatial scales (i.e., spatial frequencies) and spanning rings of different retinal eccentricities (**Figure 1**; a paradigm validated in previous work^36^). In control participants, we expect the tuning of disparity sensitivity to shift from fine to coarse spatial scales as eccentricity increases from fovea to periphery, as shown in prior studies^36,37^. On the other hand, we envision two different potential scenarios from glaucomatous vision: if glaucomatous damage is predominantly peripheral, we expect impaired disparity sensitivity in peripheral locations and low spatial frequencies; if instead glaucomatous damage involves both foveal and peripheral vision, we expect a uniform loss in disparity sensitivity across visual field locations and spatial frequencies.

**Figure 1.**
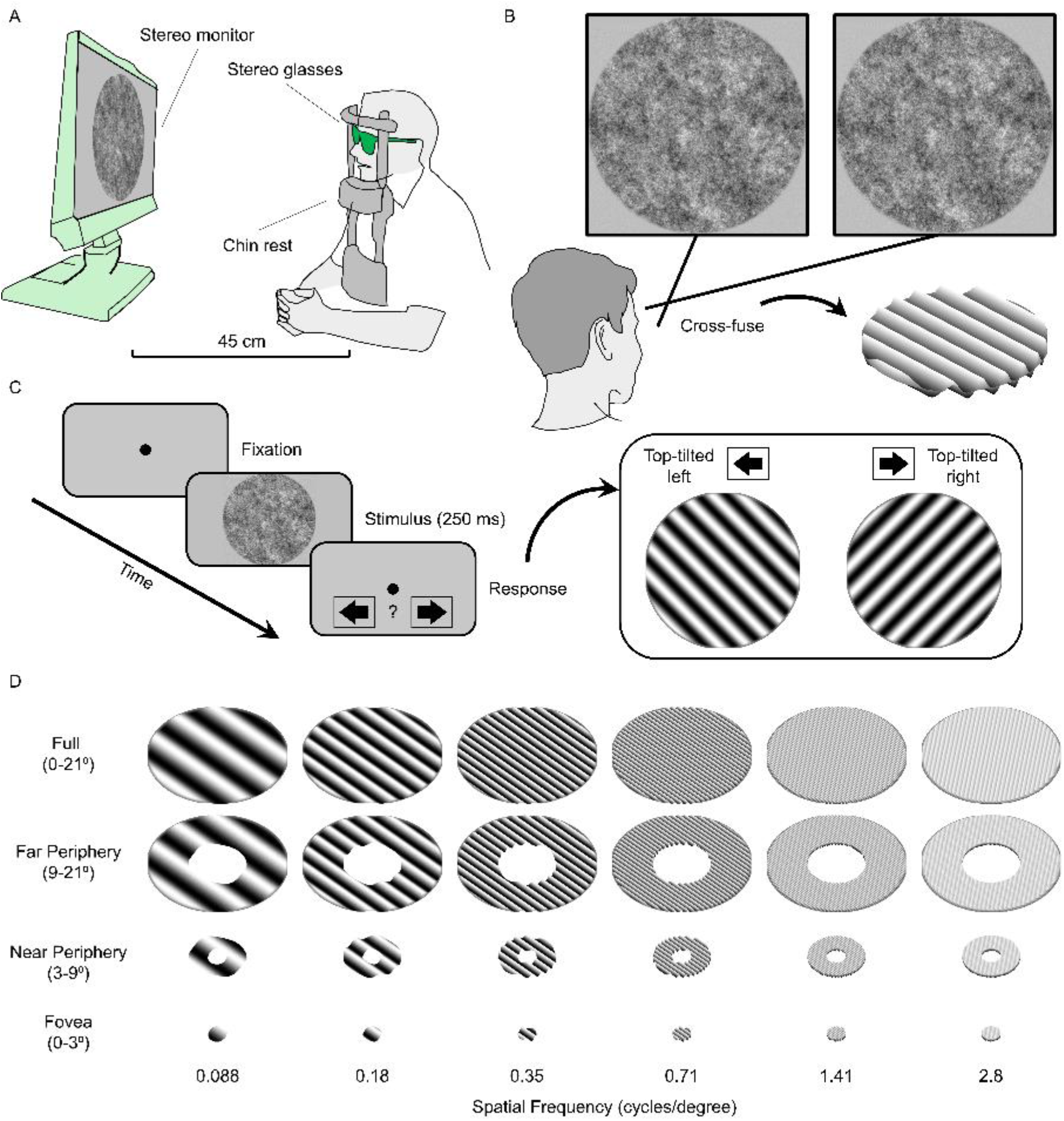
Assessing spatial frequency dependent disparity sensitivity across the visual field. **(A)** Participants seated in front of a stereo monitor viewed dichoptic stimuli through stereo shutter glasses. **(B)** Stimuli were sinusoidal disparity corrugations embedded in pink noise. Cross fuse the example stimulus pair to view the embedded 3D stimulus. **(C)** On each trial, a stimulus was shown for 250 ms, and participants were asked to report whether the disparity corrugation was top-tilted leftwards or rightwards. **(D)** Stimuli spanned 6 spatial frequencies and 4 visual field conditions.

To address the second goal, we employed the same data to adjudicate between different potential disparity integration mechanisms. To this end, the disparity sensitivity of each annular condition—representing the sensitivity of each visual field sector—was pitted against that of the full-field condition representing the integrated disparity across the visual field. Specifically, if the visual system is able to appropriately estimate and employ the reliability of binocular disparity signals at different visual field locations, then sensitivity for full-field stimuli should be equal to or better than sensitivity for stimuli spanning smaller areas of the visual field. If instead cortical integration mechanisms fails, we expect performance in the full-field condition to be worse than in the annular conditions. Depending on the extent to which cortical integration mechanisms are altered following glaucomatous damage, different scenarios can thus be speculated and compared^41,42^ as follows:

1. **Random Selection**: glaucomatous damage may impair cortical integration mechanisms to the point that the system samples disparity information from different visual field locations at random. In this worst-case scenario, disparity sensitivity to full-field stimuli would be much worse than the best sensitivity across visual field locations.
2. **Sub-optimal Integration**: glaucomatous damage may impair reliability estimates but not cortical integration mechanisms. Without being able to distinguish between visual field locations, disparity signals from all visual field locations would be averaged with equal weight. In this scenario, disparity sensitivity to full-field stimuli would be better than in the random selection case, but would still be worse than the best sensitivity across visual field locations.
3. **Optimal Selection**: the visual system may be able to estimate the ordinal reliability of disparity signals from different portions of the visual field, meaning which visual field locations contain the most reliable disparity information. If such ordinal information is available, the system could select disparity information from the most reliable visual field regions and discard the rest. In this scenario, disparity sensitivity for full-field stimuli would be equal to the best sensitivity across visual field locations.
4. **(Near-)Optimal Integration**: the visual system may be able to quantitatively estimate the relative reliability of disparity signals from different regions of the visual field. Disparity information from different visual field locations could thus be weighted by these reliability estimates and combined. If such estimates were accurate then integration would be optimal (according to the maximum-likelihood estimation principle^43–47^), and disparity sensitivity would reach its upper bound. In the more likely scenario that reliability estimates are approximate, i.e. near-optimal integration^36,48^, disparity sensitivity to full-field stimuli would nevertheless be better than the best sensitivity across visual field locations.

In sum, to evaluate the effect of glaucomatous damage and normal aging on the spatial frequency tuning of disparity sensitivity and on the way disparity is integrated across the visual field, we compare outcome measurements between patients with mild to severe glaucoma, age-matched normal controls, and young normal controls. The results of the current study shed light on the nature and inner workings of binocular integration in those with asymmetric visual field loss.

## Methods

### Participants

We recruited a total of 47 participants for this study: 18 patients with Primary Open-Angle Glaucoma (mean age 64 ± 6 years, 7 female), 16 age-matched normally-sighted older adults (mean age 61 ± 7 years, 9 female), and 13 normally-sighted young adults (mean age 25 ± 4 years, 7 female). Study participants were recruited from either Callahan Eye Hospital Clinics at the University of Alabama at Birmingham (UAB) or the UAB campus. All participants had no known cognitive or neurological impairments, further confirmed by the Mini Mental State Examination (≥ 25 MMSE score, for those aged 65 and over). Participants were fitted with proper refractive correction for the viewing distance throughout the study. Patients with glaucoma, whose diagnosis was validated through medical records, met the following inclusion criteria:

i. Glaucoma specific changes of optic nerve or nerve fiber layer defect: the presence of the glaucomatous optic nerve was defined by masked review of optic nerve head photos by glaucoma specialists using previously published criteria^49^.
ii. Glaucoma specific visual field defect: a value of Glaucoma Hemifield Test from the Humphrey Field Analyzer (HFA) outside normal limits.
iii. No history of other ocular or neurological disease or surgery that caused visual field loss.

All experimental protocols followed the tenets of the sixth revision of the Declaration of Helsinki (2008) and were approved by the Internal Review Board at UAB. We obtained written informed consent from all participants prior to the experiment, after having explained to them the nature of the study.

### Clinical assessments of binocular visual function

We assessed binocular visual function in patients and controls through standard clinical measures of binocular visual acuity (Early Treatment Diabetic Retinopathy Study charts), binocular contrast sensitivity (Pelli-Robson charts), and stereoacuity (Titmus Fly SO-001 StereoTest). We report visual acuity in logarithm of the minimum angle of resolution (logMAR), contrast sensitivity in log units (logCS), and stereoacuity in seconds of arc (arcsec). We further assessed visual field sensitivities in both eyes (24-2 test with a Humphrey Field Analyzer; Carl Zeiss Meditec, Inc., Jena, Germany) and recorded the Mean Deviation (MD) value obtained from the HFA 24-2 test, which is commonly used for evaluating the severity of glaucoma. **Table 1** reports the characteristics of study participants. According to the Hodapp—Anderson—Parish glaucoma grading system^50^, the majority of our patients were in early to moderate stages of glaucoma (13 out of 18).

**Table 1.** Characteristics of study participants.

| Diagnosis | Age<br>(Years) | Sex | Binocular<br>Visual<br>Acuity<br>(logMAR) | Binocular<br>Contrast<br>Sensitivity (logCS) | Stereoaucuity<br>(arcmin) | HFA 24-2<br>Mean Deviation (dB) |  |
| --- | --- | --- | --- | --- | --- | --- | --- |
|  |  |  |  |  |  | OD | OS |
| POAG | 62 | f | 0.02 | 1.5 | 80 | -15.58 | -6.51 |
| POAG | 63 | m | 0.02 | 1.65 | 40 | -0.61 | -2.39 |
| POAG | 68 | f | -0.08 | N/A | 40 | -0.28 | 0.81 |
| POAG | 74 | f | 0.04 | 1.65 | 40 | -1.91 | -15 |
| POAG | 62 | m | 0.12 | 1.65 | 100 | -9.55 | -10.63 |
| POAG | 67 | m | -0.1 | 1.95 | 40 | 1.54 | 0.58 |
| POAG | 56 | m | -0.02 | 1.5 | 50 | -20.91 | -27.88 |
| POAG | 73 | m | -0.1 | 1.35 | 400 | -23.65 | -23.9 |
| POAG | 65 | m | 0.02 | 1.65 | 50 | -2.38 | -15.66 |
| POAG | 64 | m | 0 | 1.95 | 40 | -0.71 | -1.71 |
| POAG | 73 | m | 0.06 | 1.65 | 50 | -5.94 | -6.94 |
| POAG | 59 | f | 0.04 | 1.65 | 50 | -1.06 | -1.75 |
| POAG | 60 | f | -0.12 | 1.9 | 40 | 1.16 | -0.55 |
| POAG | 62 | m | -0.1 | 1.9 | 40 | -2.42 | -4.64 |
| POAG | 68 | m | 0.04 | 1.7 | 100 | 0 | -7.63 |
| POAG | 59 | f | 0.04 | 2.1 | 40 | -0.3 | 1.01 |
| POAG | 55 | f | -0.06 | 1.75 | 40 | -4.25 | -7.7 |
| POAG | 60 | m | 0.04 | 1.7 | 100 | -1.02 | -6.57 |
| AMC | 51 | m | -0.2 | 2.1 | 40 | 1.6 | 0.99 |
| AMC | 55 | m | -0.18 | 1.95 | 40 | 1.98 | 1.99 |
| AMC | 65 | m | -0.12 | 1.95 | 40 | 0.74 | -0.87 |
| AMC | 63 | m | -0.08 | 1.95 | 50 | 0 | -0.24 |
| AMC | 61 | m | -0.2 | 2.1 | 40 | -1.58 | 1.35 |
| AMC | 65 | m | -0.04 | 1.95 | 100 | -0.08 | -0.73 |
| AMC | 63 | f | -0.02 | 1.95 | 40 | -1.11 | 0.01 |
| AMC | 67 | f | 0.02 | 1.95 | 40 | -1.3 | -1.75 |
| AMC | 70 | f | -0.02 | 1.95 | 60 | -0.39 | -2.41 |
| AMC | 56 | f | -0.08 | 1.9 | 60 | 3.12 | 2.05 |
| AMC | 62 | f | -0.14 | 1.95 | 40 | 2.6 | 1.07 |
| AMC | 50 | f | -0.02 | 1.95 | 40 | -0.93 | 0.59 |
| AMC | 69 | f | -0.06 | 1.95 | 40 | -0.13 | 1.59 |
| AMC | 51 | f | -0.2 | 2.1 | 40 | 0.31 | 1.83 |
| AMC | 65 | m | 0.02 | 1.8 | 50 | -1.14 | 0.9 |
| AMC | 57 | f | -0.18 | 1.95 | 40 | 2.15 | 1.06 |
| YC | 27 | m | -0.18 | 1.9 | 40 | -0.81 | -0.85 |
| YC | 25 | f | -0.22 | 2.1 | 40 | 1.02 | 1.2 |
| YC | 31 | f | -0.12 | 2.15 | 40 | N/A | N/A |
| YC | 23 | f | -0.18 | 2 | 40 | 0.19 | -1.25 |
| YC | 26 | m | -0.24 | 2.25 | 40 | 0.87 | 1.16 |
| YC | 24 | f | -0.2 | 2.25 | 40 | N/A | N/A |
| YC | 34 | m | -0.18 | 1.95 | 40 | 1.06 | 0.4 |
| YC | 26 | m | -0.06 | 1.95 | 40 | 0.35 | -0.17 |
| YC | 25 | f | -0.2 | 2.1 | 50 | N/A | N/A |
| YC | 22 | f | -0.12 | 1.95 | 40 | 1.01 | -1.58 |
| YC | 20 | m | -0.016 | 1.95 | 40 | -2.12 | -1.89 |
| YC | 20 | f | -0.2 | 1.8 | 40 | -0.25 | -0.9 |
| YC | 21 | m | -0.02 | 1.95 | 40 | -0.53 | -1.62 |

Note that POAG: primary open-angle glaucoma; AMC: age-matched control; YC: young control; f: female; m: male; OD: right eye; OS: left eye; N/A: not available.

### Measuring disparity sensitivity

We adopted a previously published experimental paradigm^36^ to measure the spatial frequency tuning of disparity sensitivity across different regions of the visual field.

#### Stimuli and apparatus

Stimuli and experimental procedures were generated and controlled using MATLAB (version 8.3; The MathWorks, Inc., Natick, MA) and Psychophysics Toolbox extensions (version 3)^51,52^ in Windows 7, running on a PC desktop computer (model: Dell Precision Tower 5810). Stimuli were presented on a liquid crystal display monitor (model: Asus VG278HE; refresh rate: 144 Hz; resolution: 1920 × 1080; dot pitch 0.311 mm) with the mean luminance of the monitor at 159 cd/m^2^. The luminance of the display monitor was linearized using an 8-bit lookup table in conjunction with photometric readings from a Minolta LS-110 luminance meter (Konica Minolta, Inc., Tokyo, Japan). Participants were seated in a dimly lit room, 45 cm in front of the monitor with their heads stabilized in a chin and forehead rest and wore active stereoscopic shutter-glasses (NVIDIA 3DVision) to control dichoptic stimulus presentation (**Figure 1A**). The cross talk of the dichoptic system was 1% measured with a Spectrascan 6500 photometer (Photo Research, Chatsworth, CA, USA). Stimuli were 1/f pink noise stereograms presented on a uniformly grey background (**Figure 1B**). Stimuli were presented as disks or rings with 1° cosinusoidal edges, and contained oblique (±45° from vertical) sinusoidal disparity corrugations of varying amplitude and spatial frequency (generated as in^36,53^, see also^54^). The central fixation target was a 0.25° black disk with 0.125° cosinusoidal edge.

#### Procedure

On each trial (**Figure 1C**), observers were presented with a black fixation dot on a uniformly grey background. As soon as the response from the previous trial had been recorded, the stimulus for the current trial was shown for 250 ms. This was too brief a presentation time for participants to benefit from changes in fixation, since stimulus-driven saccade latencies are typically greater than 200 ms^55^, saccade durations range from 20 to 200 ms^56^, and visual sensitivity is reduced during and after a saccade^57,58^. Once a stimulus had been extinguished, participants were asked to indicate, via button press, whether the disparity corrugation was top-tilted leftwards or rightwards. Participants were given unlimited time to respond, and the following trial commenced as soon as a response was provided. On each trial, we modulated the amount of peak-to-trough disparity under the control of a three-down, one-up staircase^59^ that adjusted the disparity magnitude to a level that produced 79% correct responses.

#### Design

We measured how each participant’s disparity sensitivity (1/disparity threshold) varied, as a function of the spatial frequency of the sinusoidal disparity corrugation, throughout different portions of the visual field. Specifically, we measured disparity thresholds at 6 spatial frequencies (0.088, 0.18, 0.35, 0.71, 1.41, 2.8 cycles/degree) and across 4 visual field conditions (**Figure 1D**). In the full visual field condition, stimuli were presented within a disk with a 21° radius centred at fixation. In the far and near peripheral visual field conditions, stimuli were presented within rings spanning 9°-21° and 3°-9° into the visual periphery, respectively. Finally, in a foveal condition, stimuli were presented within a disk with a 3° radius. We measured disparity thresholds for each combination of spatial frequency and visual field condition via 24 randomly interleaved staircases^59^. We combined the raw data from 50 trials from each staircase and fitted these data to a cumulative normal function via weighted least squares regression (in which the data are weighted by their binomial standard deviation). We then computed disparity discrimination thresholds from the 75% correct point of the fitted psychometric functions. We converted thresholds into disparity sensitivity following the relationship: sensitivity = 1/threshold.

#### Spatial-frequency tuning across the visual field

Disparity sensitivity is known to vary lawfully as a function of spatial frequency following an inverted-U shape^60,61^ that is well captured by a three-parameter log-parabola Disparity Sensitivity Function (DSF) model defined as^36,53^:

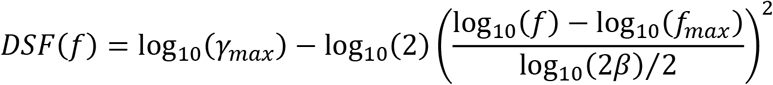

In this equation, *γ_max_* represents the peak gain (i.e. peak sensitivity), *f_max_* is the peak frequency (i.e. the spatial frequency at which the peak gain occurs), and *β* is the bandwidth at half height (in octaves) of the function. We thus fit the disparity sensitivity data to this equation separately for each visual field condition, obtaining parameter estimates that we then compared across visual field conditions and participant groups. In the full-field condition, we further computed the area under the log DSF (AULDSF) as an additional estimate of binocular function across participants.

#### Disparity integration models

Following established theories and formulations of sensory cue integration^41^, we defined four possible models of disparity integration across the visual field.

1. **Random Selection** in which disparity information is sampled from different visual field locations at random. To model this worst-case scenario, disparity thresholds to full-field stimuli were estimated from the restricted visual field conditions as:

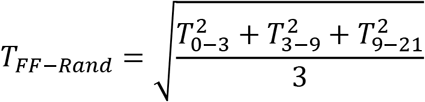
2. **Sub-optimal Integration** in which disparity information is averaged from all visual field locations with equal weight. To model this scenario, disparity thresholds to full-field stimuli were estimated as:

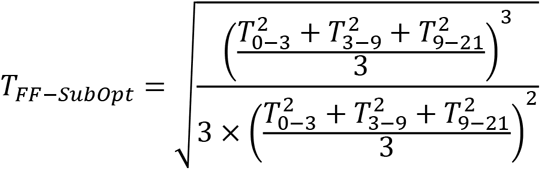
3. **Optimal Selection** in which disparity information is sampled only from the most reliable region of the visual field. To model this scenario, disparity thresholds to full-field stimuli were estimated as:

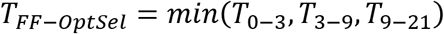
4. **Optimal Integration** in which disparity signals from different visual field locations are averaged, weighted by their relative reliability. To model this scenario, which represents the theoretical upper-bound of performance, disparity thresholds to full-field stimuli were estimated as:

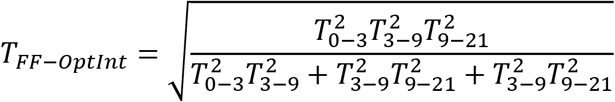

### Statistical Analyses

#### Sample Size Selection

Comparisons of binocular visual function between glaucoma patients and controls typically yield very large effect sizes (e.g.^33^, Cohen’s d>2). Effects sizes for within participant effects of interest are similarly large (e.g.^36^, Cohen’s d>1). Given that we were interested in detecting substantial effects of potential clinical significance, we exceeded a minimum sample size of N=10 per participant group. This ensured we would surpass 80% power at the 0.05 significance level for detecting effect sizes of d=1 for both between and within group comparisons. We report effect sizes for all comparisons performed in the study in terms of either Cohen’s d or η^2^, as appropriate.

#### Between-group comparisons of binocular visual function

We expected binocular visual function to be worse in old compared to young adults, and in glaucoma patients compared to both young and old healthy controls. We tested for these expected differences using one-tailed t-tests (for normally distributed data) or Wilcoxon rank-sum tests (for skewed data).

#### Comparison of spatial-frequency tuning across participant groups and visual field conditions

To test whether disparity tuning to spatial frequency varied across the participants’ visual field and across participant groups, we analysed DSF parameter estimates from the restricted visual field conditions using a 3 (participant group, between subjects factor) x 3 (visual field condition, within subject factor) mixed model ANOVA. ANOVA normality assumptions were verified via Quantile-Quantile plots. When appropriate, we conducted post-hoc comparisons via two-tailed t-tests.

#### Model selection

To adjudicate which candidate disparity integration model best accounted for the full-field data at each spatial frequency and in each participant group, we employed a simple model selection rule. Specifically, we selected as best-fitting model the one that minimized the root-mean-square error to the full-field data. If the optimal integration model won, we further confirmed whether full-field sensitivities significantly differed from the optimal selection model, using one-tailed t-tests. This was necessary to validate that full-field sensitivity truly reflected near-optimal integration. We excluded from these analyses participants whose median disparity thresholds across spatial frequency and visual field conditions were greater than 10 minutes of arc. Above this threshold, participants are either stereo blind or are not reliably performing the task^36^, and we reasoned that it would be uninformative to assess disparity integration in these participants. Based on this criterion, we thus excluded 4 glaucoma patients, 1 age matched control participant, and 1 young control participant.

## Results

### Glaucoma and normal aging exhibit expected patterns of binocular visual impairment

We evaluated binocular visual function in patients and controls using standard clinical assessment tools as described in the Methods. As expected, binocular visual acuity (**Figure 2A**) was worse in old compared to young participants (t(27)=1.9, p=.035; d=.7), and in glaucoma patients compared to both young (t(29)=5.4, p<0.001; d=2) and age-matched (t(32)=3.3, p=.0011; d=1.1) control participants. Binocular contrast sensitivity (**Figure 2B**) was not significantly different between old and young control participants (t(27)=1.4, p=.082; d=.53), but was significantly impaired in glaucoma patients compared to both young (t(28)=4.9, p<.001; d=1.8) and age-matched controls (t(31)=4.8, p<.001; d=1.7). Stereoacuity (**Figure 2C**) was significantly worse in glaucoma patients compared to young (Z=2.5, p=.0064; d=0.52) but not age-matched controls (Z=-1.2, p=.12; d=.43), and the difference between old and young control participants did not reach statistical significance (Z=-1.6, p=.057; d=.57). Finally, we visualized the mean deviation values from the HFA 24-2 test in participants’ better versus worse eyes (**Figure 2D**). The difference in HFA 24-2 mean deviation across the two eyes was much greater in glaucoma patients compared to both young (t(26)=3.5, p<.001; d=1.4) and age matched controls (t(32)=-2.7, p=.0050; d=.94), whereas young and old control participants exhibited similar interocular differences in HFA 24-2 mean deviation (t(24)=0.56, p=.29; d=.23). Together, these results clearly indicate that glaucoma patients exhibited substantial binocular visual impairment that was due specifically to asymmetric patterns of visual field loss across the two eyes.

**Figure 2.**
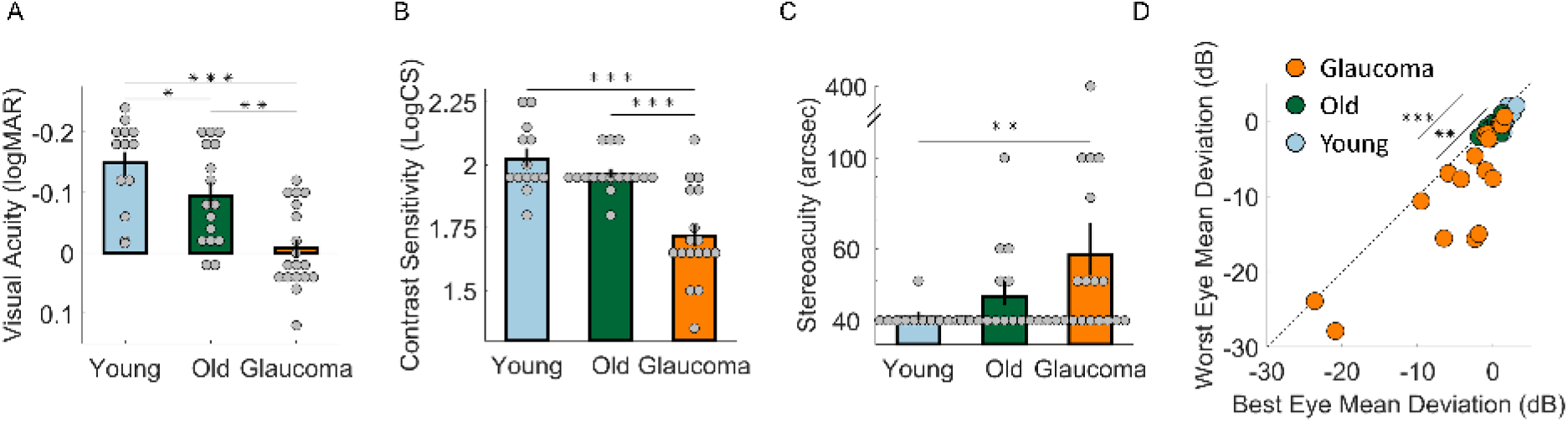
Binocular visual function in glaucoma patients and control participants. Binocular visual acuity (**A**), binocular contrast sensitivity (**B**), and stereoacuity (**C**) in patients and controls. (**D**) Scatter plot of Humphrey 24-2 visual field mean deviation in better versus worse eye. The black dotted line indicates the identity line where the MD of better eye is equal to that of worse eye. Across panels, bars are means, error bars represent bootstrapped standard error of the mean, and dots represent data from individual participants. *p<0.5; **p<0.01; ***p<0.001

### Glaucoma exhibits uniform loss in disparity sensitivity across spatial frequencies and visual field locations

Having verified that our patient cohort exhibited interocularly asymmetric glaucomatous visual field loss, we proceeded to test how this impacted disparity processing. When tested with full-field stimuli extending from the fovea to 21° into the visual periphery (**Figure 2A**), glaucoma patients exhibited a uniform loss of disparity sensitivity across spatial frequencies compared to control participants, suggesting glaucomatous damage is implicated in both the central and peripheral visual field. Indeed, the area under the log Disparity Sensitivity Function (AULDSF) fitted to the full-field condition (**Figure 2E**) was significantly reduced in glaucoma patients compared to both young (t(29)=2.2, p=.018; d=.8) and age-matched (t(32)=1.7, p=.048; d=.59) controls, whereas sensitivity did not significantly differ between young and old control groups (t(27)=0.81, p=0.21; d=.3). Further, when tested with stimuli spanning restricted portions of the visual field (**Figure 2B-D**), glaucoma patients exhibited a uniform loss of disparity sensitivity also across the visual field. In all three participant groups, the spatial frequency tuning of disparity sensitivity varied similarly across different regions of the visual field. As expected, disparity sensitivity in the far periphery (diamond markers) was tuned to depth variations at low spatial frequencies, disparity sensitivity in the near periphery (square markers) was tuned to mid spatial frequencies, and disparity sensitivity at the fovea (circle markers) was tuned to high spatial frequencies. **Figure 2F-H** further summarizes these shifts across participant groups. Specifically, the peak frequency of the disparity sensitivity curves (**Figure 2F**) shifted from high to low frequencies from the fovea to the peripheral visual field (visual field main effect: F_2,88_=260, p<.001; η^2^=.64). ANOVA results further revealed a main effect of participant group (F_2,44_=3.5, p=.039; η^2^=.033), but no interaction between visual field and participant group factors (F_4,88_=1.2, p=.31; η^2^=.006). Post hoc tests revealed a uniform shift towards higher spatial frequencies in glaucoma patients compared to young (t(29)=2.6, p=.014; d=.95) but not age-matched controls (t(32)=.63, p=.53; d=.22), and no significant difference between young and old participants (t(27)=1.8, p=.076; d=.69). The peak gain of the disparity sensitivity curves (**Figure 2G**) also decreased—as expected—from the fovea to the peripheral visual field (visual field main effect: F_2,88_=28, p<.001; η^2^=.097) and varied across participant groups (F_2,44_=7.1, p=.0021; η^2^=.18), uniformly (visual field x participant group interaction effect: F_4,88_=.35, p=.84; η^2^=.0024). Post hoc tests revealed that glaucoma patients had significantly reduced peak gain compared to both young (t(29)=3.7, p=.001; d=1.3) and age-matched controls (t(32)=2.1, p=.047; d=.71), whereas the difference between old and young participants was not statistically significant (t(27)=1.8, p=.082; d=.67). Finally, the bandwidth of disparity tuning (**Figure 2H**) remained constant across visual field locations and patient groups (visual field main effect: F_2,88_=.11, p=.9; η^2^=.001; participant group main effect: F_2,44_=2.7, p=.081; η^2^=.061; visual field × participant group interaction effect: F_4,88_=.88, p=.48; η^2^=.017). These patterns confirmed previous reports regarding the tuning of human disparity sensitivity across different regions of the visual field^36^. More importantly, and contrary to the commonly held belief that early glaucomatous damage is predominantly peripheral, these results demonstrated that glaucoma patients suffered a uniform loss of disparity sensitivity across both foveal and peripheral visual sectors.

### All groups exhibit near-optimal disparity integration across the visual field

Our results thus far determined that interocularly asymmetric glaucomatous visual field loss leads to impaired disparity sensitivity both foveally and peripherally, and across spatial frequencies. Does this in turn impair the way disparity signals are integrated across the visual field? To test this, we used the sensitivity data from the restricted visual field conditions to generate predictions for full-filed sensitivities in four possible scenarios: random selection, sub-optimal integration, optimal selection, and optimal integration, as described earlier. Across all three participant groups (**Figure 3I-K**) and across nearly all spatial frequencies, full-field sensitivity data approached the upper bounds of possible performance and was best fit by either the optimal selection or optimal integration models. Furthermore, full-field sensitivities were significantly better than the optimal selection model—and thus conclusively reflected near-optimal integration—predominantly at low spatial frequencies. Specifically, disparity integration was significantly near-optimal at the lowest spatial frequency tested in young participants (**Figure 3I**, 0.088 cycles/degree: t(11)=2.6, p=.013; d=.74), age-matched controls (**Figure 3J**, 0.088 cycles/degree: t(14)=3.1, p= .0039; d=.8), and glaucoma patients (**Figure 3H**, 0.088 cycles/degree: t(13)=4, p<.001; d=1.1). Performance was also significantly near optimal at the second lowest spatial frequency tested in both age-matched controls (**Figure 3J**, 0.18 cycles/degree: t(14)=2.4, p=.015; d=.62), and glaucoma patients (**Figure 3H**, 0.18 cycles/degree: t(13)=6.1, p<.001; d=1.6). These results indicate that participants reliably integrated low-spatial frequency disparity information most clearly across far- and mid-peripheral visual field sectors. At higher spatial frequencies instead, participants may have relied more dominantly on foveal disparity estimates. Critically, even though glaucoma patients exhibited significant asymmetric visual field defects and impairments in disparity processing, cortical mechanism for disparity integration appeared to be spared.

**Figure 3.**
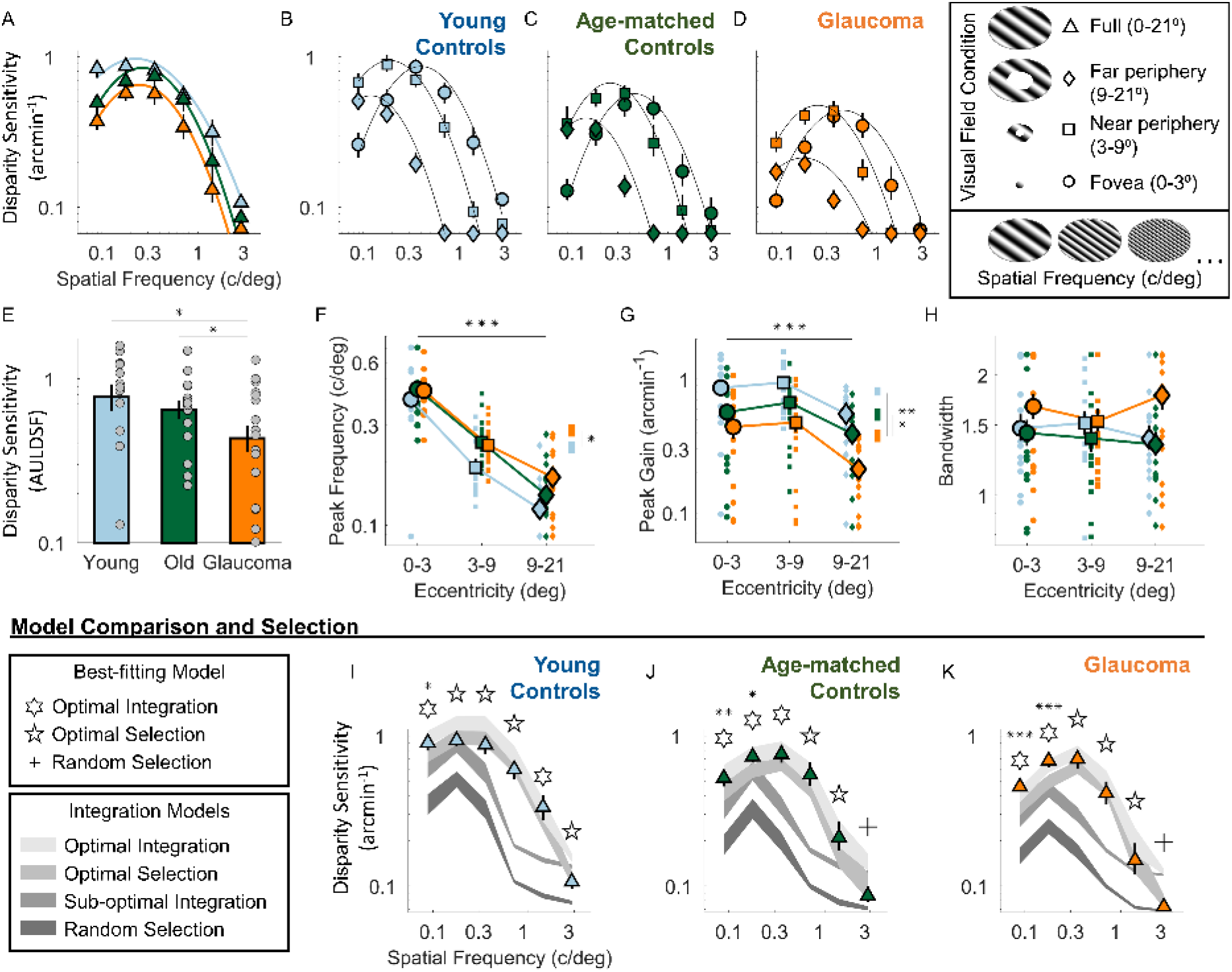
Disparity sensitivity across the visual field and integration model selection. (**A**) Disparity sensitivity plotted as a function of spatial frequency for full-field stimuli in patients and controls. (**B-D**) Disparity sensitivity as a function of spatial frequency for stimuli spanning far (diamonds), near (squares), and foveal (circles) portions of the visual field, plotted separately for young controls (B), age-matched controls (C), and glaucoma patients (D). In (A-D), continuous lines are the average best fitting log parabola functions passing through the data. (**E**) Area under the log Disparity Sensitivity Function for full-field stimuli in patients and controls. (**F-H**) Peak frequency (F), peak gain (G), and bandwidth (H) of the fitted log parabola models as a function of the portion of visual field tested, in patients and controls. (**I-H**) Model predictions and psychophysical measurements of disparity sensitivity for full-field stimuli, plotted as a function of spatial frequency for young controls (I), age-matched controls (J), and glaucoma patients (K). Across panels, bars and large markers are means, dots and small markers represent data from individual participants, error bars and shaded regions represent bootstrapped standard error of the mean. *p<0.5; **p<0.01; ***p<0.001

## Discussion

Binocular disparity is a key component of depth perception. In healthy humans, foveal regions preferentially process disparities at fine spatial scales, peripheral visual regions are tuned for coarse spatial scales, and the visual cortex selects and combines depth information across different visual regions by accounting for these differences in tuning^36^. Glaucoma is a neurodegenerative condition characterized by progressive loss of retinal ganglion cells and resulting visual field defects. Even in early stages of the disease, glaucomatous ganglion cell damage results in patterns of visual sensitivity loss that may vary both across the visual field and between the eyes^15,19,27,62^. Additionally, damage to ganglion cells can potentially propagate to cortical regions due to direct and transsynaptic anterograde axonal degeneration^38–40^ For this reason, it is highly plausible that glaucoma could lead to impairments in binocular disparity processing across the visual field and even along the cortical visual processing pathway. Thus, here we assessed how glaucomatous visual field damage impacted spatial-frequency dependent disparity sensitivity across different sectors of the visual field.

Using a previously validated experimental paradigm^36^, we assessed the spatial-frequency dependent disparity sensitivity across the visual field and further determined the best integration model accounting for the full-field disparity sensitivity data of glaucomatous vision. Our results demonstrate several fundamental aspects of glaucomatous visual loss. First, we observed a uniform loss in disparity sensitivity across visual field locations and spatial frequencies in glaucoma, compared to both young and age matched healthy controls. This demonstrates that foveal vision and binocular function are impacted even in early-glaucoma. Second, we found that disparity sensitivity to full-field stimuli was equal to or better than sensitivity for stimuli spanning smaller areas of the visual field, in glaucoma and control patients alike. The glaucomatous visual system is thus able to access—at least to some extent—the reliability of the signals coming from different regions of its damaged visual field. Further, early glaucomatous loss does not substantially impact cortical selection and integration mechanisms, leading to near-optimal processing of binocular disparity even given the presence of binocularly asymmetric glaucomatous lesions across the visual field.

Previous studies have reported that binocular function, including stereopsis, is significantly impaired in glaucomatous vision^27,33^. However, glaucomatous stereovision deficits have been characterized almost exclusively in terms of stereoacuity, and such deficits have been identified even in glaucoma suspects^35^. This is perhaps surprising, given how glaucomatous damage is believed to spare the central vision until the end stages of the disease, whereas stereoacuity refers to fine spatial scales that should be processed at the fovea. Our results reconcile this apparent contradiction by demonstrating that glaucoma patients exhibit a loss of disparity sensitivity across spatial scales, from the fovea to the visual periphery. Furthermore, our findings lend strength to the view that macular damage may commonly occur even in early stages of glaucoma^14–21^.

The observed glaucomatous deficits in stereovision across the visual field could have affected cortical processing in several ways. In the most extreme scenario, glaucomatous neurodegeneration could have propagated from retinal regions upwards along the visual pathway^38–40^, reaching cortical regions responsible for selecting and combining disparity depth information across the visual field. Alternatively, even if cortical integration mechanisms had been spared, glaucomatous damage could have unpredictably altered the reliability of disparity signals across the visual field, rendering integration processes sub-optimal. Instead, our analyses revealed that glaucoma patients performed near optimal integration of disparity information across visual field sectors, particularly at coarse spatial scales preferentially processed across peripheral visual regions. This finding is far from trivial, since patients with glaucoma are often unaware of the localization of their visual field deficits, particularly when these are asymmetric between the eyes^63,64^. It is thus notable that the visual system can appropriately select disparity information from different visual field sectors and even combine this information, weighted by the relative reliability of the disparity signals. This suggests not only that cortical disparity integration mechanisms are spared, but that—at least in early or moderate stages of the disease—the system is able to adapt and make the best possible use of the remaining binocular visual function. Our findings are indeed consistent with previous work^65^ demonstrating that the mechanism underlying the binocular summation of contrast sensitivity (i.e., a quadratic summation rule with an exponent of 1.3) remains well preserved even in early and moderate glaucoma patients. Taken together, cortical binocular integration mechanisms, whether the signal is luminance contrast or disparity, appear to be spared in glaucomatous vision despite binocularly asymmetric vision loss.

While such findings hint at potential treatment opportunities for preserving or restoring binocular function^32,66–70^ in early or moderate glaucoma, we acknowledge that whether this would occur also in more advanced disease stages needs to be addressed in future studies. In addition, more quantitative assessments^25,36,71,72^ of the relationship between the pattern of glaucomatous visual field defects and the pattern of impairment in disparity processing would help us further characterize binocular disparity processing throughout the visual field.

## Acknowledgements

The authors would like to thank Marguerite Devereux, Rong Liu, and Lindsay Washington for their assistance with subject recruitment and data collection. This research was supported by the DFG (German Research Foundation: project No. 222641018-SFB/TRR 135 TP C1), NIH/NEI Grant R01 EY027857, and Research to Prevent Blindness (RPB) / Lions’ Clubs International Foundation (LCIF) Low Vision Research Award.

## Data availability

Data and analysis scripts will be made are available from the Zenodo database upon acceptance.

## Author contributions

G.M. and M.K. conceived and designed the study. M.K. collected the data. G.M. performed the analyses. G.M. and M.K. wrote the manuscript.

